# Autocatalytic–Protection for an Unknown Locus CRISPR–Cas Countermeasure for Undesired Mutagenic Chain Reactions

**DOI:** 10.1101/2020.03.24.004291

**Authors:** Ethan Schonfeld, Elan Schonfeld, Dan Schonfeld

## Abstract

The mutagenic chain reaction (MCR) is a genetic tool to use a CRISPR–Cas construct to introduce a homing endonuclease, allowing gene drive to influence whole populations in a minimal number of generations^1,2,3^. The question arises: if an active genetic terror event is released into a population, could we prevent the total spread of the undesired allele^4^? Thus far, MCR protection methods require knowledge of the terror locus^5^. Here we introduce a novel approach, an *autocatalytic-Protection for an Unknown Locus* (a-PUL), whose aim is to spread through a population and arrest and decrease an active terror event’s spread without any prior knowledge of the terror-modified locus, thus allowing later natural selection and ERACR drives to restore the normal locus^6^. a-PUL, using a mutagenic chain reaction, includes (i) a segment encoding a non-Cas9 endonuclease capable of homology-directed repair suggested as Type II endonuclease Cpf1 (Cas12a), (ii) a ubiquitously-expressed gene encoding a gRNA (gRNA1) with a U4AU4 3′-overhang specific to Cpf1 and with crRNA specific to some desired genomic sequence of non-coding DNA, (iii) a ubiquitously-expressed gene encoding two gRNAs (gRNA2/gRNA3) both with tracrRNA specific to Cas9 and crRNA specific to two distinct sites of the Cas9 locus, and (iv) homology arms flanking the Cpf1/gRNA1/gRNA2/gRNA3 cassette that are identical to the region surrounding the target cut directed by gRNA1^7^. We demonstrate the proof-of-concept and efficacy of our protection construct through a Graphical Markov model and computer simulation.

## Main

A directed CRISPR–Cas9 mutagenic chain reaction (MCR) in the germ line of a sexually reproducing animal, where homology-directed repair (HDR) inserts a construct containing (i) a segment encoding Cas9 (ii) a ubiquitously-expressed gene encoding a gRNA specific to some desired genomic sequence and (iii) homology arms enclosing the Cas9/gRNA insert that are identical to the region surrounding the target cut directed by the encoded gRNA, has been shown to generate an autocatalytic spread of this gene drive through the population even if introducing a negative fitness^2,6^. An MCR gene drive, having already worked in mammals, has enormous potential, including the ability to regulate and even eradicate the spread of malaria by female *Anopheles* mosquitoes^2^. However, what happens if in the process, an unintended genetic modification is introduced by the gene drive to an entire species? Moreover, the simplicity of the MCR design and its introduction to an organism’s germ line creates the potential for active genetic terror^8^. A gene encoding a tumor suppressor, such as p53, can be knocked out of an entire population by introducing an MCR with gRNA targeting the gene in a fraction of the population and allowing the autocatalytic reaction to spread the gene drive (carrying a null mutation) to near 100% of progeny with each successive generation^9^. In a quickly reproducing species such as *Anopheles*, generating high offspring counts, and even higher when mated in laboratory, a terror event could result in the non-lethal ubiquitous loss of a protein in the population. The possibility extends to the same effect in a sheltered small human population. The same transformative innovation that MCR brings to the control of diseases, is just as great in its ability to catalyze genetic terror. With the advent of MCR, it is essential to find a prompt means of curtailing the potential spread of an Undesired MCR through a population.

The driven gene is inherited by each generation, through HDR, with a probability ‘*h*’ of approximately 0.95<*h*<1.0. Non-homologous end joining (NHEJ), can lead to an evolved-resistant allele to the homing endonuclease^6,10^. Because NHEJ occurs at a non-zero probability, 1-*h*, at each generation, the gene drive will lose its ability to spread in a population over some number of generations. By only relying on nuclease-induced resistant mutations, the frequency of animals with the driven gene is 20% after 25 generations according to previous models^9^. Thus, in an active genetic terror event, using this stand-alone method will likely not confer resistance to a population at a reasonable time frame.

Engineered strategies for reversing a CRISPR–Cas9 MCR spread of a driven gene and its homing endonuclease include synthetic resistance, reversal drives, immunized reversal drives, ERACR, and e-CHACR constructs^5,6,11^. A discrete-time model for these measures has shown varying successes, largely dependent on the number of animals originally introduced with the Undesired drive and the number of animals introduced with the countermeasure. Effective countermeasures, such as immunizing reversal drives, target both the Undesired driven gene and the wild-type population, conferring resistance. Other effective countermeasures such as e-CHACR (a second site “construct hitchhiking on the autocatalytic chain reaction”), target the homing endonuclease Cas9, blocking autocatalytic ability^6,11^. Frequency-only population genetic models show that about 20 generations are required for a small 1:10 release of a synthetic resistant allele countermeasure to bring the Undesired driven allele frequency to <0.1; about 15 generations are required for a small 1:10 release of a reversal drive countermeasure, and about 10 generations are required for a large 1:1 release of an immunizing reversal drive countermeasure^5^. However, these measures either require prior knowledge of the specific loci targeted by the Undesired drive, which would not be available in a terror event until large-scale population RNAseq and genomic analysis could be performed, or require the release of the countermeasure drive only after a majority of the population has the Undesired drive.

We introduce a countermeasure construct, an *autocatalytic-Protection for an Unknown Locus* (a-PUL), that does not require prior knowledge of the locus modified by the Undesired MCR and does not require the majority spread of the Undesired driven allele prior to countermeasure release (Figure 1). This construct, like e-CHACR, will make a double-strand break in the Cas9 homing endonuclease, and like immunizing reversal drives, will use its own homing endonuclease to drive itself through a population. a-PUL includes (i) a segment encoding a non-Cas9 endonuclease capable of homology-directed repair of a double-strand break such as Type II endonuclease Cpf1 (Cas12a), (ii) a ubiquitously-expressed gene encoding a gRNA (gRNA1) with a U4AU4 3′-overhang specific to Cpf1 and with crRNA specific to some desired genomic sequence of non-coding DNA, (iii) a ubiquitously-expressed gene encoding two gRNAs (gRNA2/gRNA3) both with tracrRNA specific to Cas9 and crRNA specific to two distinct sites of the Cas9 locus, and (iv) homology arms flanking the Cpf1/gRNA1/gRNA2/gRNA3 cassette that are identical to the region surrounding the target cut directed by gRNA1.

**Fig. 1:**
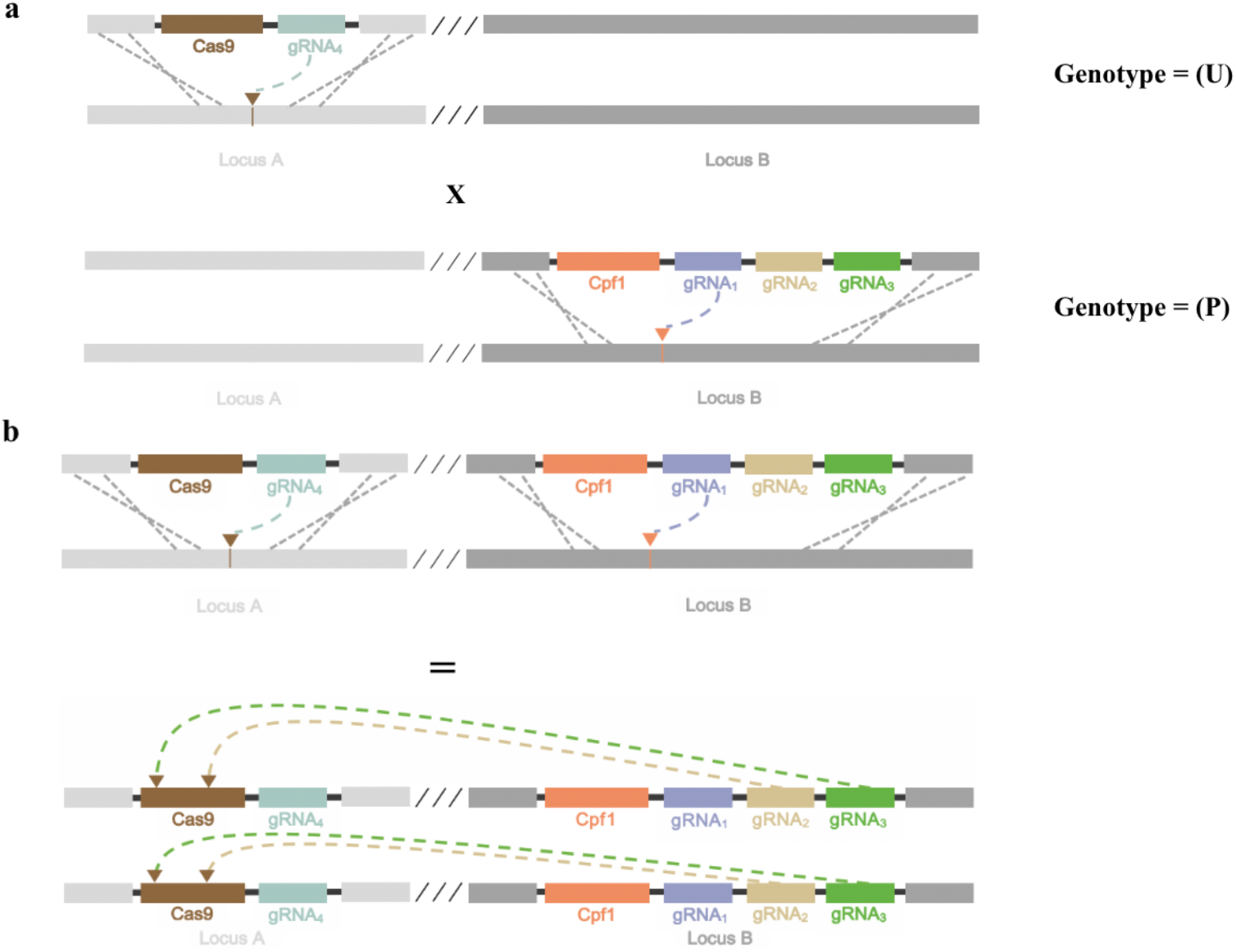
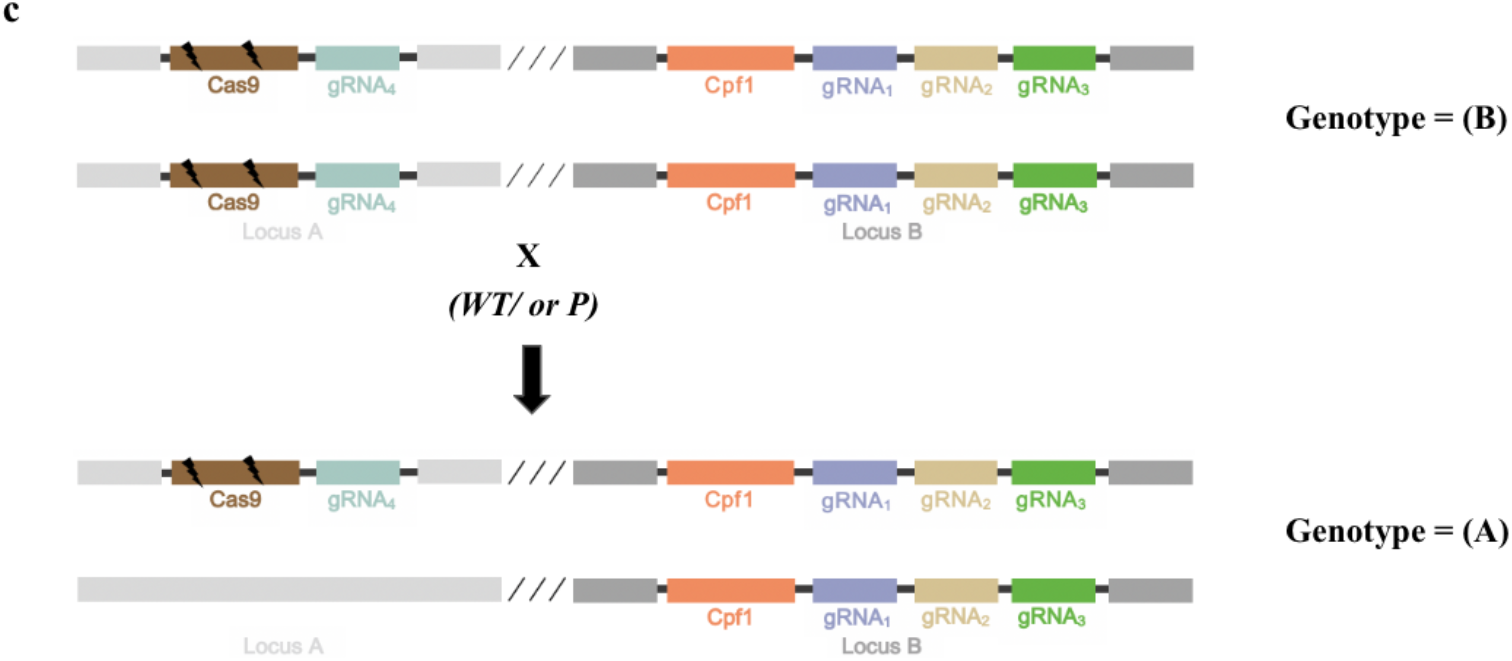
A cross between an Undesired MCR (U) and a-PUL (P) with F1 crossed to WT/or P restoring wild-type allele. **a**, An Undesired MCR with wild-type allele at Locus B, represented by the genotype (U), targets Locus A, creating a null mutation. An animal with the a-PUL countermeasure inserted into a non-coding DNA region and wild-type allele at locus A, represented by the genotype (P) is crossed with (U). Both will use a distinct homing endonuclease to drive the desired allele into the homologous chromosome and will pass down an identical allele to the F1. Figure 1 shows results assuming HDR (0.95<*h*<1.0). **b**, The F1 inherits the driven gene from both the (U) and (P) parents. The Cas9 homing endonuclease serves both to drive the construct from (U), as well as to form a CRISPR–Cas complex with gRNA2 and gRNA3 to induce a double-strand break and eventual NHEJ with endonuclease derived mutation in the homing endonuclease from (U). Thus, both alleles of Locus A of the F1 will be unable to undergo future MCR while the a-PUL protection construct will continue to drive. **c**, After NHEJ in both Cas9 alleles of Locus A, Locus A carries a biallelic-null mutation, but has no gene drive ability, and Locus B has a-PUL. We will call this exact genotype (B), shown at the top of Fig. 1.c. Crossing of the F1, genotype (B), with wild-type, genotype (W), or with genotype (P), will always result in F2 with the (A) genotype shown at the bottom of Fig. 1.c. Thus we demonstrate that for parental U and P and mating F1’s with either W or P will result in a restored wild-type allele at the locus targeted by the Undesired MCR, as well as inhibiting any future MCR of the Undesired MCR by creating a double mutation in its homing endonuclease.

Cpf1 is a novel class of smaller CRISPR–Cas endonucleases. After inducing a double-strand-break at the target site, Cpf1 leaves a short overhang end which is more conducive for precise DNA sequence insertion^12^. Both LbCpf1 and AsCpf1, derived from *Lachnospiraceae* and *Acidaminococcus*, respectively, using 1000-bp long arms, induced a donor cassette through HDR^13^. LbCpf1 increases HDR in zebrafish, suggesting potentially increased a-PUL efficacy compared to an MCR utilizing Cas9^14^. Because NHEJ can introduce resistant mutants to the homing endonuclease, if HDR frequency is improved in the a-PUL relative to the Undesired MCR, the a-PUL countermeasure can overtake the Undesired MCR in fewer generations than other countermeasures.

To determine the efficacy of such a construct, we developed a Graphical Markov model to measure the failure rate, the population frequency of biallelic-null mutation, of the proposed approach by deriving the genotype probability corresponding to each potential genotype possible from Figure 1. Because NHEJ can induce resistant mutations to the Undesired MCR over generations, we adopt a conservative model that ignores the decrease of the Undesired MCR due to NHEJ derived mutations in the model^10^. Therefore, if the computational simulation, ignoring nuclease-induced mutations, of the models succeed, we gain even greater confidence in the proposed a-PUL countermeasure.

To derive a term for genotype probability in an offspring of a given generation, we show all possible matings of genotypes for each of the first three generations (Figure 2). We then show that all possible matings of genotypes for each subsequent generation is identical to the transition from the second to third generation. W animals carry the wild-type genotype, P carry the a-PUL construct at Locus B, U carry the Undesired MCR genotype, B carry the genotype shown on Figure 1 resultant from a cross of U and P, and A carry the genotype shown on Figure 1 resultant from a cross of B and W.

**Fig. 2:**
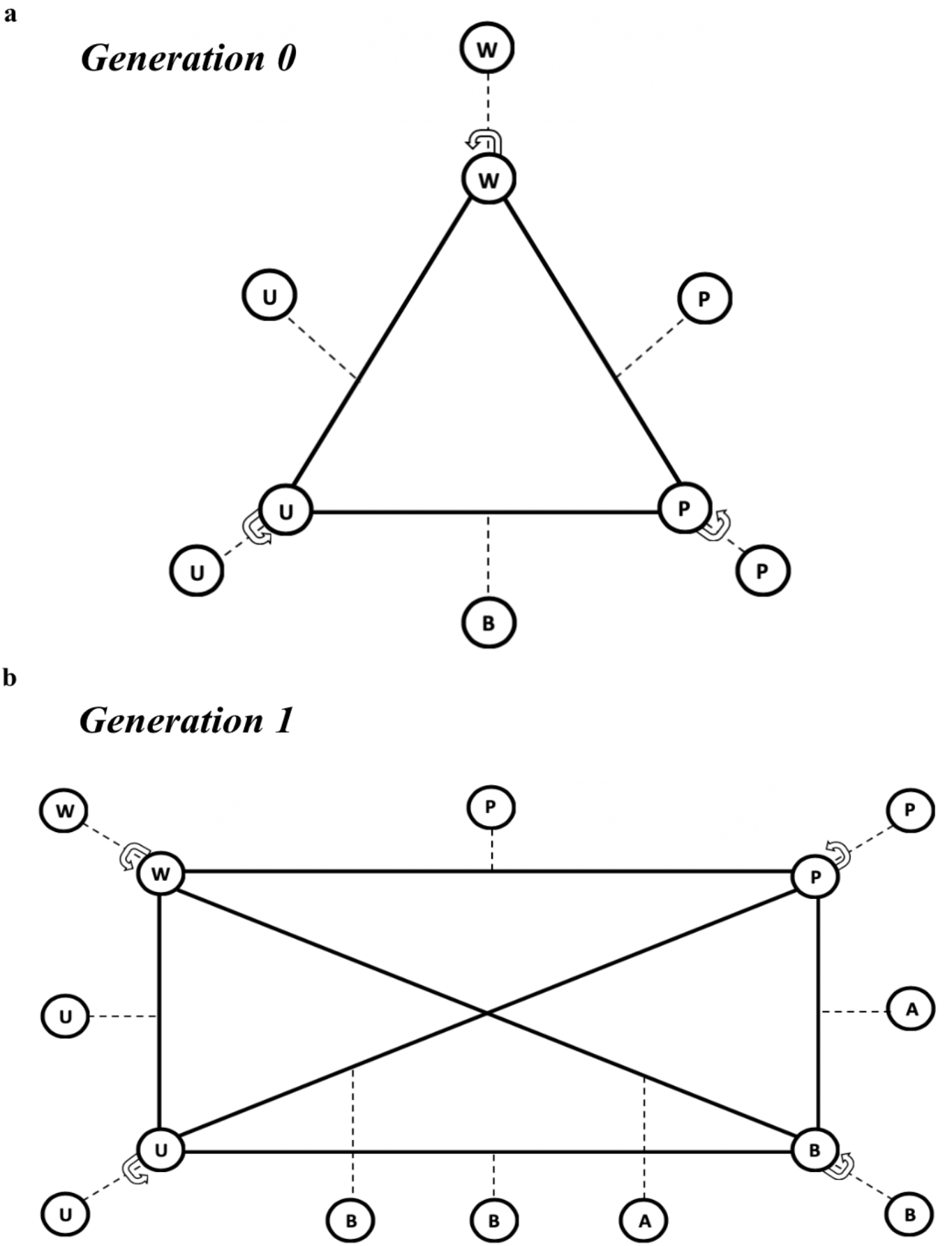
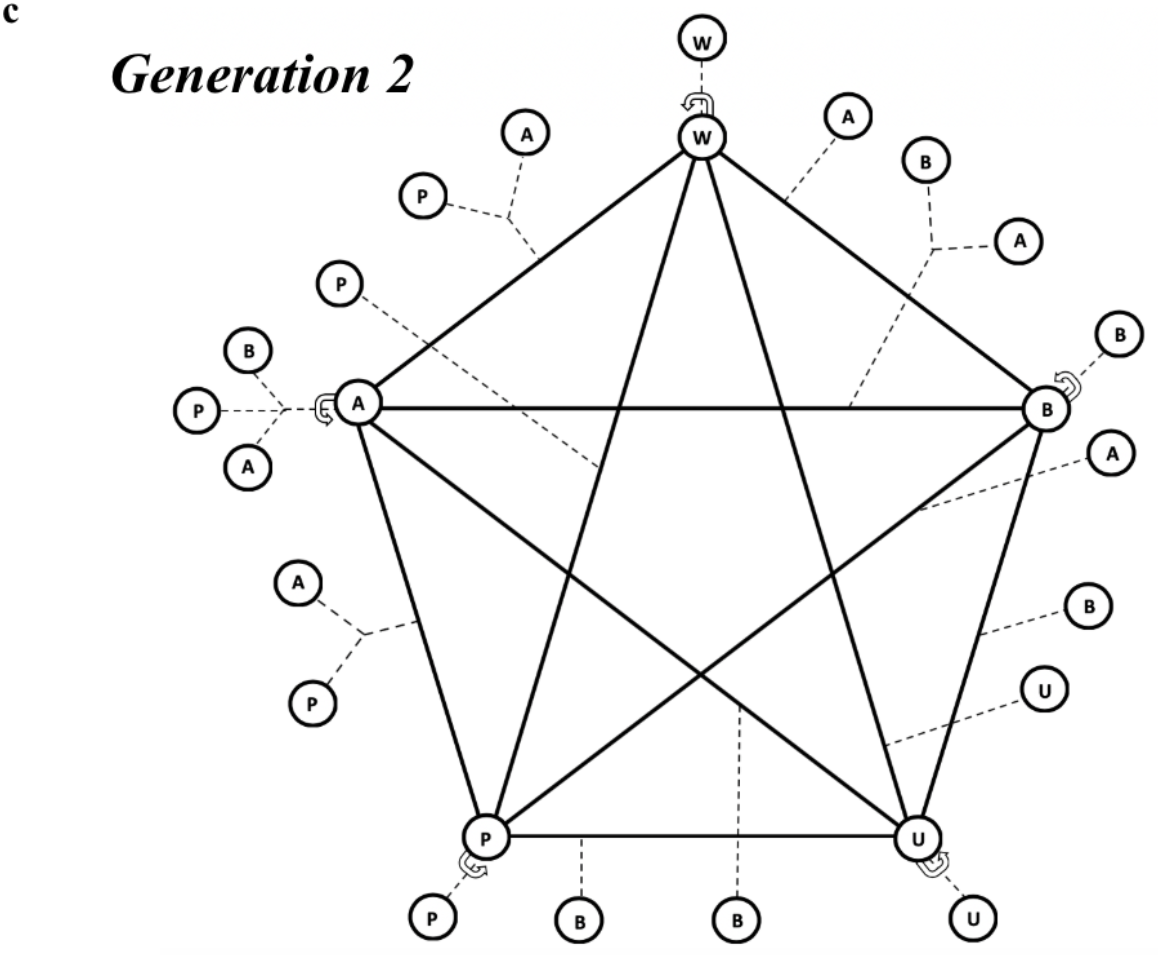
Constructing a probability map by generation for all possible genotype matings to define a recursive model. **a**, Generation 0, the parental generation of the model, begins with a wild-type population with genotype (W) with some animals released who are of (U) genotype, carrying the Undesired MCR. There are also some animals added who are of (P) genotype, with the a-PUL Gene Drive. All matings, assuming HDR, are shown either by filled in black lines or a circular arrow for the same genotype mating. Dashed lines represent offspring genotypes. Mating *generation 0* animals gives rise to a new genotype (B) (see Figure 1). **b**, Mating *generation 1* animals gives rise to a new genotype (A) (see Figure 1). **c**, The fraction of genotype frequencies when multiple genotypes branch from the same mating is determined by simple Mendelian inheritance; When A is mated to A, B will arise at ¼ frequency, P at ¼ frequency, and A at ½ frequency. Similarly, when A is mated to B, A will arise at ½ frequency and B at ½ frequency. Mating *generation 2* animals gives rise to no new genotypes. Therefore, genotype frequency in the population can be recursively modeled starting at *generation 3*, based on the frequencies of genotypes (W), (U), (P), (A), (B) in *generations 0-2*.

We present here the number of wild-type animals in the population w, the number of Undesired MCR carrying animals first released into the population u, and the number of a-PUL carrying animals first released into the population p. We also present *h* as the probability of HDR after every double-strand break.

Based on Figure 2, with probabilities derived from Mendelian crosses of W, U, P, A, and B shown in Figure 1, we derive the probabilities P_k_(W), P_k_(U), P_k_(P), P_k_(A), and P_k_(B) of an offspring of *generation k* to correspond to each of the five genotypes W, U, P, A, and B, respectively, computed as the number of possible matings N_k_(W), N_k_(U), N_k_(P), N_k_(A), and N_k_(B) corresponding to each genotype, respectively, divided by the total number of possible matings M_k_ for each generation *k*.

Parents’ generation:

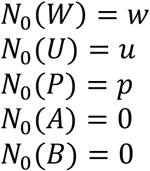

First-offspring generation:

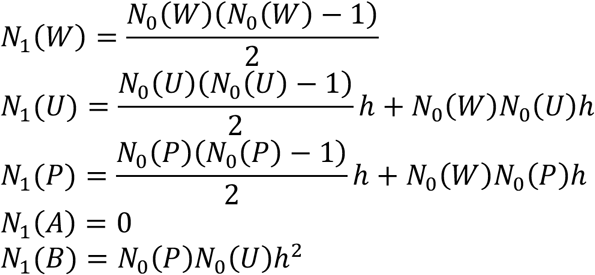

Second-offspring generation:

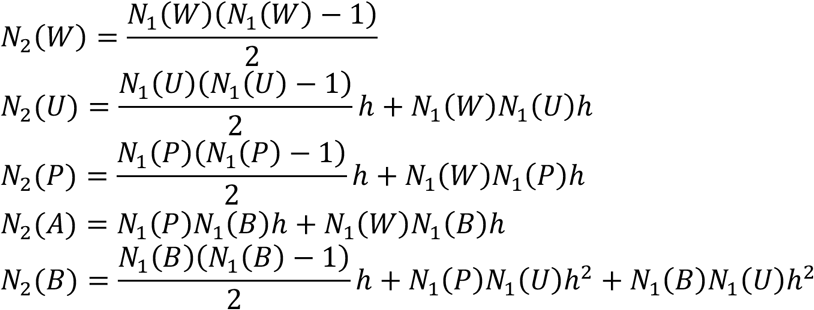

Offspring generation k ≥ 3:

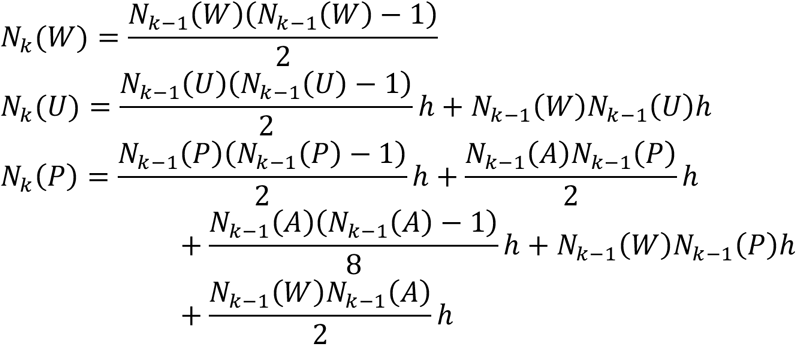

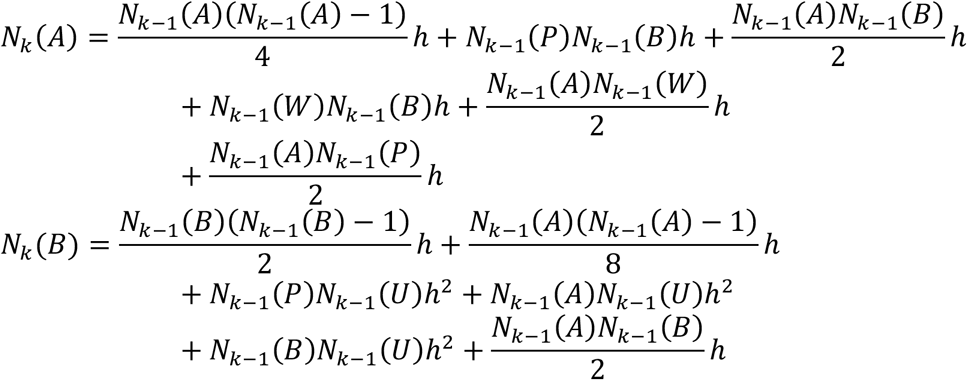

where

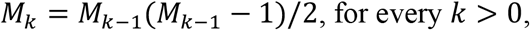

and *M*_0_ = *w* + *r* + *p*.

Finally, we note that P_k_(W), P_k_(U), P_k_(P), P_k_(A), and P_k_(B) are given by N_k_(W), N_k_(U), N_k_(P), N_k_(A), and N_k_(B) divided by M_k_, respectively, for each generation *k*; *e.g.*, P_K_(*W*) = N_K_(*W*)/*M*_K_, for every *k* ≥ 0.

Success of the proposed a-PUL countermeasure is achieved when the offspring do not carry a biallelic-null mutation on the Undesired MCR target locus; *i.e.*, offspring have genotype W, P, or A. We therefore measure the *failure rate* 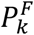 of the a-PUL countermeasure given by

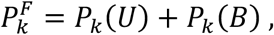

for generation *k*. The failure rate 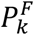 represents the probability of an offspring of generation *k* carrying a biallelic-null mutation on the Undesired MCR target locus. In this model, the Undesired MCR targeted a locus of some gene coding for a protein where one copy is sufficient for wild-type phenotype. If we can arrest probabilistic growth of the biallelic-null mutation, once the specific locus is determined, constructs such as ERACR or functions such as natural selection can reduce the frequency of these genotypes in successive generations.

Figure 3 depicts the failure rate for the genotypes above computed through seven generations and plotted against the number of generations. In Figure 3, we show several examples of successful a-PUL ability to arrest growth of biallelic-null mutation frequency in the population. Graphically, both a plateau or a decrease in the frequency of the biallelic-null mutation in the population represent success, as once the Undesired MCR is arrested its effects may be reversed by the methods discussed above.

**Fig. 3:**
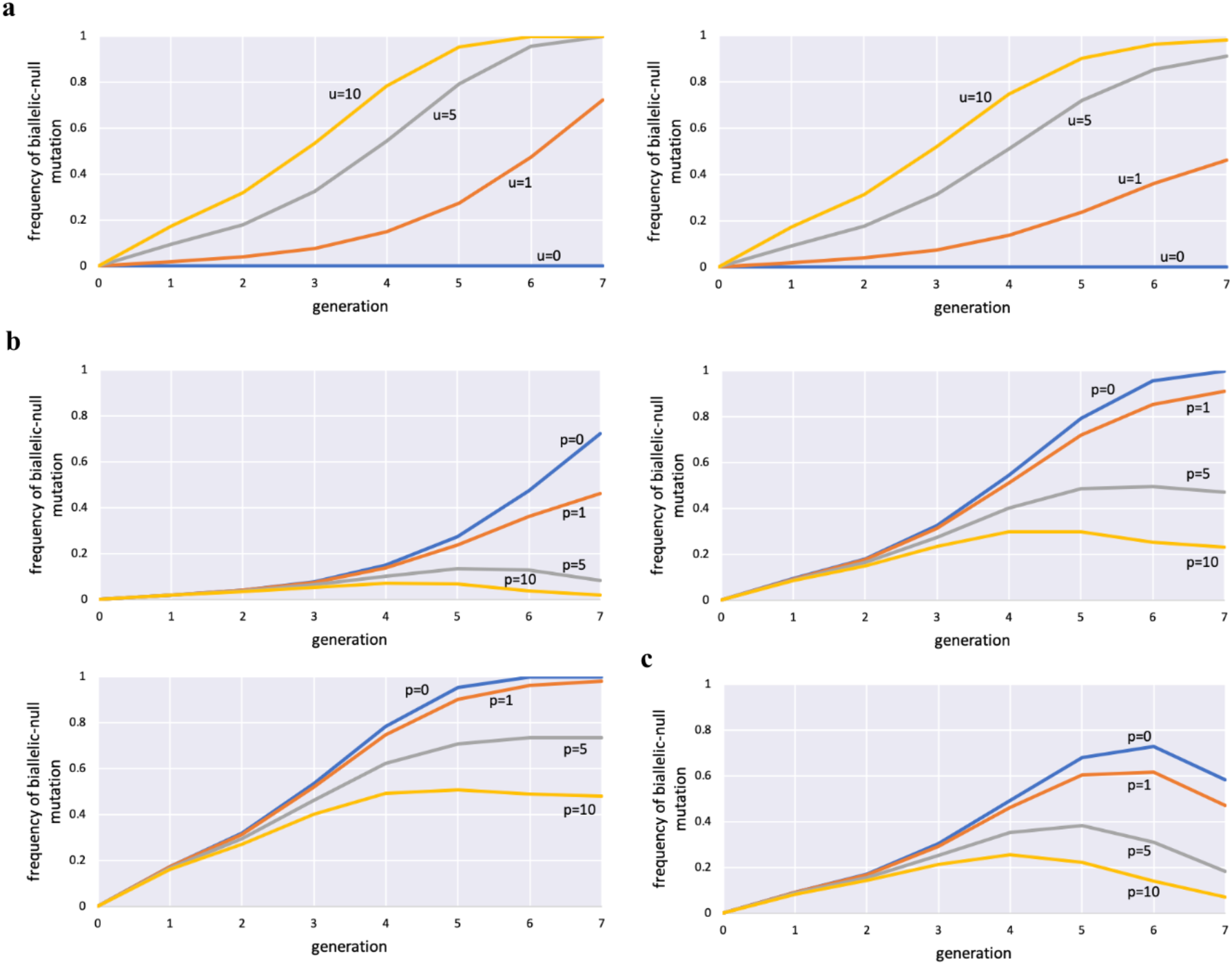
Successful a-PUL ability to arrest growth of biallelic-null mutation in a population is dependent on the number of Undesired MCR and number of a-PUL genotypes released. **a**, Wild type population is always set at 100, as this will scale with the number of Undesired MCR (u) and number of a-PUL (p). (Top left) zero a-PUL genotypes are introduced in the while the number of Undesired MCR is scaled 0, 1, 5, and 10 (w=100; u=0,1,5,10; p=0; h=1). (Top right) The same scaling of Undesired MCR is done in the right graph, however, one a-PUL genotype is introduced, arresting the spread of biallelic-null mutation (w=100; u=0,1,5,10; p=1; h=1). **b**, (Top left) An Undesired MCR is introduced and the number of a-PUL genotypes released is scaled from 0, 1, 5, 10 (w=100; u=1; p=0,1,5,10; h=1). (Top right) the same scaling of p is introduced when u=5 (w=100; u=5; p=0,1,5,10; h=1). (Bottom left) the same scaling of p is introduced when u=10, demonstrating that a 1:1 release of a-PUL to Undesired MCR can prevent spread of biallelic-null mutation (w=100; u=10; p=0,1,5,10; h=1). **c**, (Bottom right) NHEJ induced resistant mutations will eventually halt the spread of any homing endonuclease. Here we show that when probability of HDR is 0.98, the coupling of an a-PUL countermeasure to NHEJ induced resistance will result in greatly decreased biallelic-null mutation frequency in the population (w=100; u=5; p=0,1,5,10; h=0.98).

Many genetic constructs have been suggested to dilute the spread of an Undesired MCR through a population, and some success has been demonstrated. However, the success is largely dependent on prior knowledge of the specific locus targeted by the Undesired MCR, or by allowing the driven allele to have already penetrated through the majority of a population such that an endonuclease is already expressed that a countermeasure construct could use.

We have introduced a-PUL, *autocatalytic-Protection for an Unknown Locus*, which relies on using an endonuclease, capable of generating a double-strand break and allowing for HDR, that is not Cas9. By not using Cas9 in a-PUL, the gRNA on the a-PUL countermeasure can target the Cas9 DNA on the Undesired homing endonuclease and thus inhibit the ability for the Undesired drive to use MCR to spread through super-Mendelian inheritance through a population. The endonuclease on a-PUL can theoretically be any other than Cas9 that fits the above criteria; here we suggest Cpf1 for this function as Cpf1 is smaller than Cas9 and has been suggested in prior studies to allow for HDR at greater frequency^14^. LbCpf1 induces the double-strand break about 18 nt away from PAM site allowing for repeated cleavages to increase the frequency of HDR, whereas Cas9 generates base-pairing mismatches which prevent repeated cutting of the target site after an NHEJ event^14^. This allows a-PUL to spread with greater efficacy in a population by less frequently generating resistant alleles through NHEJ than the Undesired MCR generated by using Cas9. If an Undesired MCR uses a Cpf1 endonuclease, the a-PUL can be revised to include Cas9 or any other suitable endonuclease.

Future work should address the selective deactivating of the a-PUL sequence in the event that the Undesired MCR has been driven to a null frequency in subsequent generations. While gRNA2 and gRNA3 are feedforward activated by the presence of Cas9, and thus the activated CRISPR– Cas system would only be present in animals who have the Undesired MCR, gRNA1 is always capable of forming the activated CRISPR–Cas system with Cpf1. Continued formation of any CRISPR–Cas complex always carries a potential for off-target effects. Thus, novel systems such as including a CRE-LOX system encasing the a-PUL homing endonuclease could selectively slow the autocatalytic spread of a-PUL through a population after a terror event or Undesired MCR has been properly controlled^15,16^.

In summary, mutagenic chain reactions (MCR) can autocatalytically spread a desired allele through an entire population in just a number of generations. The CRISPR–Cas MCR has the potential to change animal genetics by spreading resistance to many diseases for future generations. For example, it has the potential to use a fast-reproducing system such as *Anopheles* to spread immediate changes that can even affect human lives over large regions. Just as the last decade of the CRISPR–Cas system gave us editing tools over the genome, we introduce a-PUL as a needed “undo button” as gene-drives begin to develop across species and generations.

## Methods

Computer simulations were performed using GNU Octave, version 4.4.0, and our simulation code is available upon request. To run the program, we used a 2019 MacBook Pro (13-inch), 1.4 GHz Intel Core i5 Processor, 8 GB 2133 MHz LPDDR3 memory, running macOS Mojave Version 10.14.6.

The Graphical Markov model proposed relies on the observations that in a generic genotype class *S* having *s* animals with the same genotypes, there exist 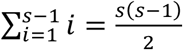 possible offspring, whereas between two generic genotype classes *S* and *T* having *s* and *t* animals, respectively, there exist *st* possible offspring. The probability of a generic genotype class, in a particular generation, is determined by the ratio of the number of possible offspring for that genotype divided by the total number of possible offspring for that generation. We note that support for the veracity of our model has been attained by having confirmed that the total sum of probabilities of all genotype classes (*i.e.*, W, U, P, A, and B) is unity for each generation.

## Author Information

*Stanford University, Stanford, CA, USA*

Ethan Schonfeld

*Multimedia Communications Laboratory, University of Illinois, Chicago, IL, USA*

Elan Schonfeld

Dan Schonfeld

### Contributions

Ethan S. and D.S conceived the project. Ethan S. designed the genetic constructs. Ethan S., Elan S., and D.S generated and analyzed the models. Elan S. performed the computer simulations. Ethan S. analyzed the simulations. Elan S. generated the figures. Ethan S. wrote the manuscript with input from all authors.

